# Selective volumetric excitation and imaging for single molecule localization microscopy in multicellular systems

**DOI:** 10.1101/2022.12.02.518828

**Authors:** Tommaso Galgani, Yasmina Fedala, Romeo Zapata, Laura Caccianini, Virgile Viasnoff, Jean-Baptiste Sibarita, Rémi Galland, Maxime Dahan, Bassam Hajj

## Abstract

Light sheet fluorescence microscopy (LSFM) has become a leading standard in high-resolution imaging of living samples in 2- and 3-dimensions. Biological samples are however not restricted to a single observation plane and several molecular processes evolve rapidly in 3D. The conventional mechanical scanning required in LSFM limits the range of observable dynamics and are usually restricted in resolution. Here we introduce a new strategy for instantaneous volumetric excitation and volumetric imaging of single-molecules in cell aggregates. The technique combines, for the first time, the use of light sheet microscopy and multifocus microscopy (MFM) and enables a volumetric 4D imaging of biological samples with single-molecule resolution. We engineered the excitation beam to yield a modular and uniform excitation matching the observable detection range of MFM. The strength of the method is highlighted with examples of single-molecule 3D tracking and 3D super-resolution imaging in multicellular samples.

## Introduction

Biological processes evolve at various temporal and spatial scales. Observing such processes in living volumetric samples and at the molecular scale remains challenging. Light-sheet fluorescence microscopy (LSFM)^1^ has become a main approach for 3D imaging of live samples over prolonged durations due to its high sectioning capability, improved contrast, low photobleaching and reduced phototoxicity. Conventional LSFM relies on focusing and fast linear teetering of a laser beam to form a thin excitation sheet located at the focus of a second collection objective. As the number of detected photons affects directly the localization precision of single molecules (SM)^2,3^, a high numerical aperture (NA) collection objective is desirable. Relying on high NA objective is however compromised by the orthogonal configuration of conventional light-sheet microscopes.

In recent years, different strategies have been implemented to adapt LSFM to high NA collection objectives. For instance, lattice light sheet microscopy circumvents the mechanical limitations by custom engineered objectives and sample holders^4^. In Tilted light sheet microscopy with 3D point spread functions (TILT3D), the excitation Gaussian beam is tilted compared to the observation focal plane^5^. This was shown to improve 3D SM imaging despite the angular mismatch. Other methods rely on a reflective surface of an AFM tip^6^ or a custom made prism for redirecting the light sheet^7^. In contrast to using two objectives, another class of LSFM methods relies on using a single-objective for exciting and imaging the sample. A first approach relies on tilting the excitation sheet similarly to HILO^8^ and introducing a two-objective remote imaging system along the emission path of the microscope. The first objective forms a primary volumetric image of the sample, while the second objective is tilted to image an inclined view of the primary image. The tilt angle between the two objectives is chosen to match the orientation of the light-sheet^9–12^. A second approach, named single-objective Selective Plane Illumination Microscopy (soSPIM), relies on placing a reflective surface near the sample to reflect the excitation beam and to form the desired sheet perpendicular to the optical axis of the observation objective^13–15^.

However, cells are three-dimensional objects and intracellular events are typically not constrained to one focal plane. Conventional light-sheet microscopy relies on a sequential scanning of the focal plane and the light-sheet through the sample. This method is inadequate for detailed studies of fast intracellular and single molecule dynamics in 3D. Indeed, the focal plane may frequently be at the wrong place at the wrong time, thereby missing important aspects of dynamic events. In addition, while the mentioned methods proved to be beneficial for studying monolayers of cultured cells, there is a lack of techniques dedicated for fast SM imaging in multicellular systems such as in cell colonies or organoids, at a distance from the coverslip.

In this paper we introduce a new modality for volumetric light sheet fluorescence microscopy (V-LSFM) dedicated to single-molecule imaging in multicellular systems. Our method achieves high temporal and spatial resolutions, provides a good signal to noise ratio (SNR) over the volume of the sample and reduces photobleaching and phototoxicity. We show how a modular and tunable system built around a standard inverted fluorescence microscope can optimally combine soSPIM^13^ and MultiFocus Microscopy (MFM)^16,17^ in order to produce the selective excitation and instantaneous observation of a volume. Our approach relies on a uniform axial excitation based on shaping the intensity profile of the excitation beam. In combination with an adapted sample holder, our method proved to be efficient in imaging SM in clusters of cells 10 to 25 *μm* away from the coverslip using a 60×/1.27 NA objective. Volumes of 1 to 4*μm* thickness are imaged at 50Hz, a frequency that is compatible with imaging fast SM dynamics. We illustrate the advantage of our method by performing single particle tracking in both synthetic samples and live cells, as well as super-resolution imaging by DNA-PAINT at high concentrations.

## Results

### Design of the selective homogeneous volumetric microscope

Following the 3D dynamics of fast diffusing single molecules or performing super-resolution imaging within cell aggregates are challenging tasks. To address these challenges, we set several requirements for the development of our efficient V-LSFM, notably that the system must (i) selectively excite individual molecules tens of microns within the sample, (ii) image an entire volume with an adjustable axial extension at an acquisition rate up to 50 Hz per volume to enable fast 3D single particles tracking, (iii) provide adjustable excitation thickness, ranging from 1 to 4 *μm*, to suit the extent and acquisition speeds of different samples and (iv) produce a uniform excitation over the observed axial range. To achieve these requirements, we primarily capitalized on two techniques: the single-objective selective plane illumination (soSPIM) and the multifocus microscopy (MFM). We specifically developed and optimized different optical modules, each one performing individual tasks yet acting jointly to achieve an efficient volumetric selective excitation and imaging (Figure 1 A).

**Figure 1:**
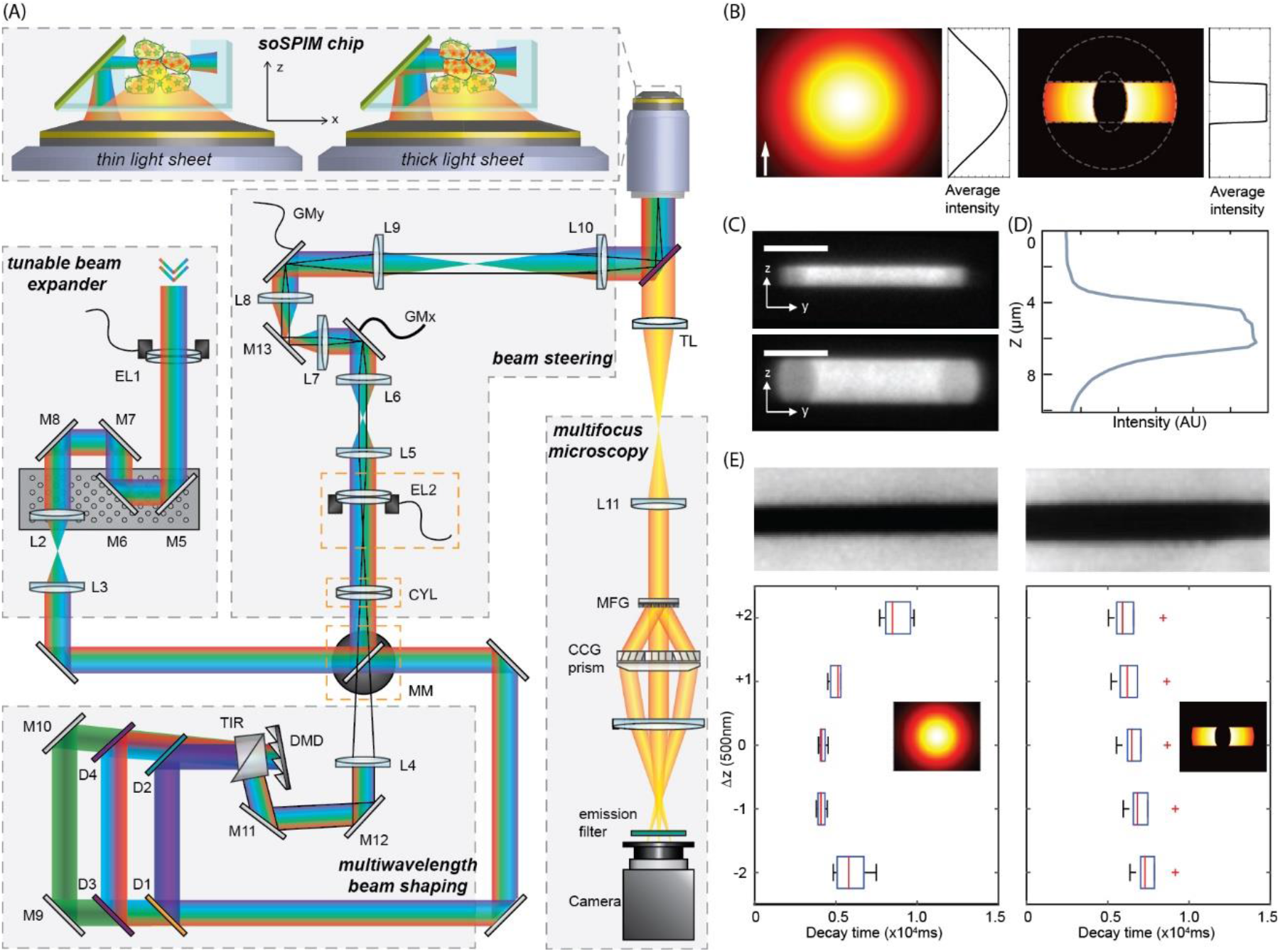
V-LSFM principle. (A) Schematic presentation of the setup. A tunable beam expander adjusts the size of the excitation beam. The beam shape is engineered using a DMD conjugated to the sample plane. A mirror placed at the sample plane reflects the excitation beam to form the light sheet. The combination of the automatized beam expander and beam shaping allow to control the thickness of the light sheet to match the observation volume of multifocus microscope (MFM) introduced at the emission path. (B) The beam shape is engineered to yield a uniform excitation over the thickness of the light sheet. A double arc beam shape yields >90% uniform and a confined excitation over the desired imaging volume. (C) Fluorescence response of the double arc beam when scanned to produce the homogeneous axial profile. The intensity profile is measured by direct projection of the light sheet over a uniform fluorescence layer without any reflection. The double arc size and profile defines the thickness of the uniform light sheet. (D) The axial fluorescence profile shows a confined excitation with >80% uniformity over the desired thickness. (E) Characterization of the photobleaching induced by scanning a gaussian beam in contrast to a double arc-shaped beam: in double arc excitation the photobleaching is uniform over the light sheet thickness.

In the soSPIM technology, cells are seeded in a microfabricated sample holder containing several arrays of microwells positioned at close proximity from a 45° mirror. With this geometry, the mirror reflects the excitation laser inside the sample, allowing creating a thin light-sheet using the same high NA objective used to collect the fluorescence. The beam steering module is composed of two galvo-mirrors and an electrical tunable lens conjugated to the back focal plane of the objective, and a third mirror conjugated to the sample plane. It allows performing the fast-scanning of the beam along the mirror axis and adjusting the optimal position and the excitation laser waist to create a light-sheet perpendicular to the optical axis of the objective and well positioned on the sample (requirement (i)). Yet, acquiring a volumetric image in soSPIM, as for most of light-sheet methods, still requires a mechanical scanning of either the sample or the objective and light-sheet position, limiting the volumetric acquisition speed and therefore the range of observable dynamics in 3D.

In MFM, multiple planes are acquired simultaneously on the same camera in a single exposure, without the need for any mechanical movement. MFM is based on the incorporation of several diffractive elements on the emission path of a microscope. Up to 9 focus-shifted and aberration-free images are formed and tiled on the same camera. Since no mechanical movement is involved, the achievable volumetric temporal resolution is only limited by the fluorescence brightness of the fluorophore and the camera frame rate. The central element in MFM is a multifocus grating (MFG) conjugated to the back focal plane of the objective. The main motif of the MFG is designed to yield a uniform light distribution between the desired diffraction orders while maximizing the diffraction efficiency at multiple imaging wavelengths^18^. To cope with the requirement of instantaneous 3D observations of adjustable volumes (requirement (ii)), different MFG motifs were designed to yield different number of diffraction orders (i.e imaging planes). We have designed different MFGs with 4 (MFG4) and 9 (MFG9) imaging planes (check supplementary information and Supplementary Figure 1). When combined with a 60× magnification objective, MFG4 and MFG9 result in an imaging volume thickness of 2 and 4*μm* respectively, as compared to the μm range depth of field of this objective. The gratings are mounted on magnetic bases for a smooth exchange, without the need of realignment. The gratings are also compatible with the same chromatic correction module.

To match the excitation and the desired observation volume, we implemented an automated beam expander module for fine tuning the illumination size of the Gaussian beam. The challenge here is to produce lightsheet thicknesses ranging from 1 to 4 *μm*. Such range requires extended magnification factors that are difficult to implement in a conventional beam expander. Our beam expander is based on the combination of three lenses: 1) an electric lens (EL) with adjustable focal lens, 2) a lens mounted on a translation stage and 3) a lens fixed on the optical table. The optical configuration is adjusted such that any displacement *s* of the translation stage would result in increasing the distance between the first and second lens by *s* and shortening the space between the last two by 3*s* (Supplementary Figure 2). Different combinations of the EL focal length and the stage displacement produce a collimated Gaussian beam demagnified by a factor ranging between 1× and 0.2×. The size of the Gaussian beam at the input of the objective determines the waist of the excitation at the focus and thus the effective light-sheet thickness. The module is completely automated using a LabVIEW program and can generate very thin light-sheets (1 *μm* thickness) for high optical sectioning over short distances (Rayleigh length of 5*μm*, Supplementary video 1), as well as thick light-sheets (up to 4 *μm* thickness) for confined volumetric excitation over large distances (Rayleigh length of 78*μm*) (Supplementary information) (requirement (iii)).

Still, achieving uniform imaging over extended volumes requires the excitation of the sample with a uniform intensity distribution over the desired axial range. A light-sheet excitation produced by fast linear teetering of a Gaussian beam is characterized by an axial intensity profile that follows a Gaussian curve. A large-size gaussian beam results in a quasi-homogeneous excitation near its central peak but extends the excitation outside of the desired observation range, thus compromising the advantages of a selective excitation. To create a selective excitation with a homogenous axial intensity, we developed a beam shaping module (requirement (iv)) based on a Digital Micromirror Device (DMD). In the DMD each pixel of a micromirror array can be switched independently between an ON (on axis reflection) and OFF (off axis reflection) states. As a result, the intensity profile of the reflected beam is shaped according to the pattern applied on the DMD. The individual pixels of the DMD act as a 2D blazed grating which implies wavelength dependent diffraction that was compensated by using a combination of multiple dichroic mirrors and prisms (Methods and Supplementary Figure 3). The DMD is placed on a plane conjugated to the focal plane of the objective, enabling the modulated shape of the excitation beam to be reproduced at the sample plane. We carried out a simulation which demonstrated that a beam imprinted with an elliptical double arc shape produces a uniform volumetric excitation when scanned laterally (Check supplementary information and Supplementary Figure 5-6). We characterized the uniformity of the excitation by imaging a homogeneous fluorescent layer illuminated with the scanned shaped beam. We found that the intensity profile was >80% homogenous over the entire volume (Figure 1 B and Supplementary Figure 9-10). The extension of the excitation volume was tuned between 2*μm* and 5*μm* by regulating the size of the incoming Gaussian beam using the beam expander module and adjusting the size of the DMD pattern accordingly. To illustrate the importance of a uniform excitation we compared the photobleaching curves obtained by illuminating a homogenous fluorescent layer (Figure 1 C-E) with a standard Gaussian light-sheet and with a double arc light-sheet. We characterized the fluorescence decay due to photobleaching at different distances from the center of the light-sheet. As expected a Gaussian light-sheet results in a non-homogeneous bleaching, with a shorter decay time at the center of the excitation compared to the edge. By contrast, the light-sheet obtained by linear teetering of the elliptical double-arc shaped beam produces homogenous photobleaching over the excitation volume as shown by the decay time of the fluorescence intensity curves (Figure 1 E).

The illumination homogeneity of the double arc beam was compared to the homogeneity obtained by the direct 2D scanning of a focused excitation beam. The approach relies on scanning a 1 *μm* thick Gaussian beam along the mirror, to excite the desired volume. In this configuration, the beam shaping module is bypassed to yield a classical focused Gaussian beam, which in turn is scanned laterally and axially by the beam steering module to produce an extended illumination. We found that the 2D scanning of a Gaussian beam can be exploited to produce >80% homogenous excitation for volumes extending over > 5 *μm*. We compared the homogeneity of the intensity distribution of such illumination to the intensity profile of the double arc shaped illumination. The latter was found more suitable for a 1*μm* to 5*μm* light-sheet thickness range (Supplementary Figure 10) while preserving a confined excitation over the full range of a cell. An additional factor to consider, is the extent of the excitation produced by the light-sheet along its propagation direction. Due to light diffraction, the light-sheet excitation remains confined over a distance proportional to the Rayleigh length from the beam waist. For a volumetric light sheet obtained by a beam shaped as a double arc, the waist can be approximated as *ω*′_0_ = N × *ω*_0_. Here the waist *ω*′_0_ is expressed as a *N* times the waist *ω*_0_ obtained for a diffraction limited focused Gaussian beam. Compared to a light-sheet of the same waist obtained by 2D scanning of a focused Gaussian beam, an excitation beam shaped as a double arc extends over 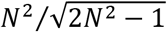 larger Rayleigh length (Supplementary Information). Based on the uniformity and the extent of the light-sheet, the double arc-shaped beam was thus chosen for generating the light-sheet in this paper.

The combination of the different modules on the excitation and emission path results in a modular system for selective homogeneous excitation and instantaneous volumetric imaging. The setup is capable of recording tunable volumes of 22μm×22 μm×4μm, ~20 *μm* away from the coverslip surface, with an imaging rate reaching 50 volumes/s. To highlight the power of achieving instantaneous volumetric imaging, we performed 3D single-particle tracking (SPT) in synthetic and biological samples. We also performed super-resolution imaging of densely labeled samples using DNA-PAINT.

### V-LSFM allows the recovery of long trajectories

As compared to 2D confined molecules, tracking fast diffusing particles in 3D volumes, such as within the nucleus of cells, require instantaneous volumetric imaging to enable the recording of long trajectories. As an illustration, we followed the dynamics of 100 nm and 200 nm fluorescent beads freely diffusing in PBS with an expected diffusion of 4 and 2*μm*^2^/*s* respectively. Thanks to our V-LSFM system, various ranges of single-particle dynamics could be explored (Protocol in Methods) (Supplementary video 2). The recorded trajectories were analyzed (check Methods) and compared to conventional single plane imaging. In the latter configuration, due to the limited depth of field of a 2D imaging system (~600 nm), the imaged particles tend to diffuse outside of the focal plane within 1*s*. As a result, short trajectories (<50 frames, 20 ms/frame) are obtained and the dynamic information is partially lost. Moreover, the same particles can appear multiple times in the focal plane which tends to increase the apparent number of trajectories. The histogram of the computed diffusion coefficients might overestimate the proportion of fast diffusing particles. By contrast, by extending the imaging depth in V-LSFM, we could track the same fluorescent beads diffusing at 3.8*μm*^2^/*s* over 6s. We tuned the acquisition volume using different MFGs and we matched the size of the homogeneous selective excitation by finely adjusting the size of the DMD pattern. We imaged volumes of 22μm×22μm×4μm without any mechanical scan. Compared to 2D imaging, the length of the recorded trajectories was found to outspread over longer time scales, up to 300 frames, acquiring 50 vol/s (Figure 2 A and Supplementary video 3).

**Figure 2:**
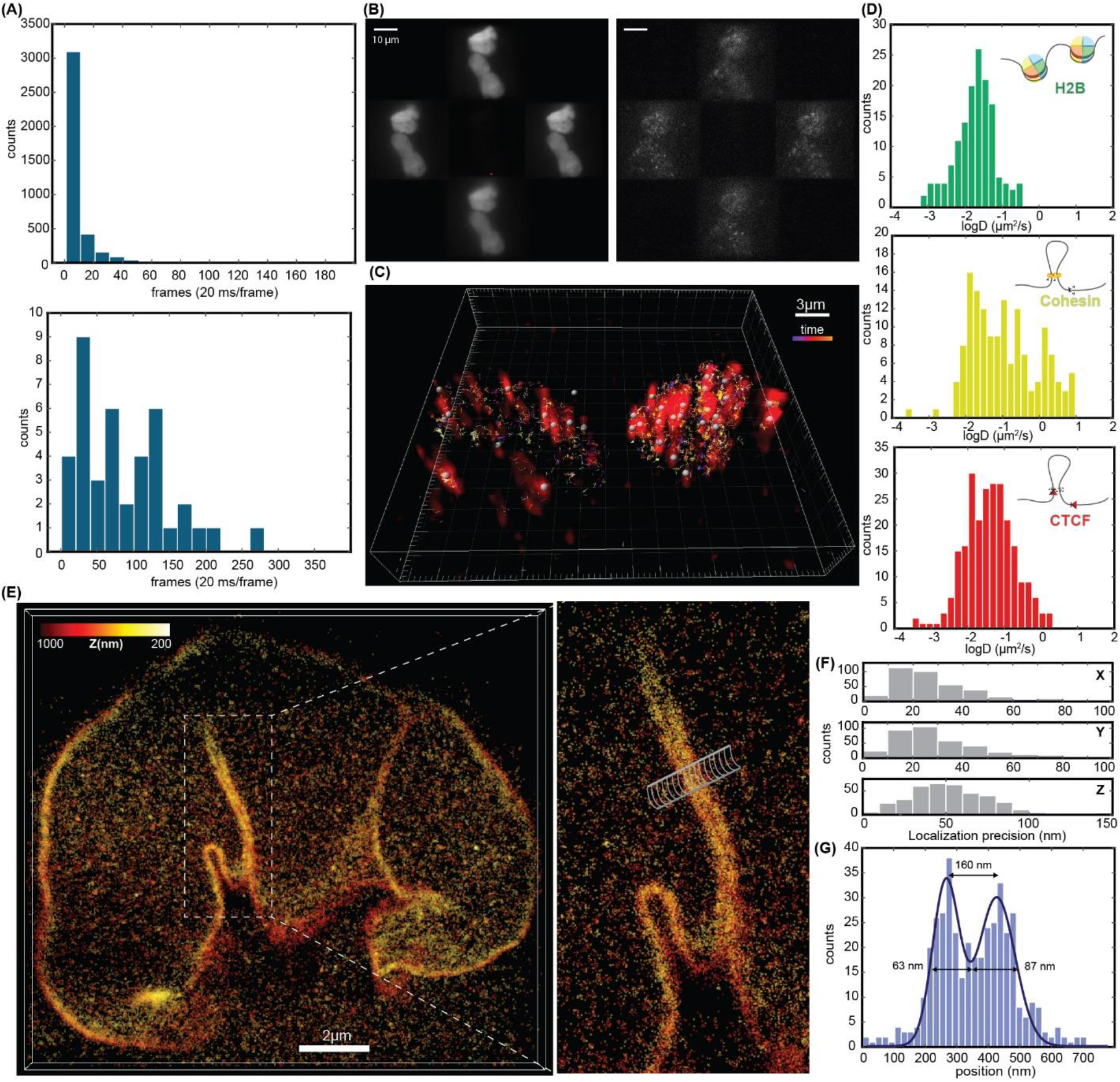
Single molecule localization microscopy using V-LSFM for studying the dynamics and organization at the single molecule level. (A) Histograms of the recovered trajectory length when imaging particles in V-LSFM compared to 2D: Tracking single particles in V-LSFM increases trajectories lengths compared to 2D tracking which over-estimates the number of trajectories. (B) Example of live cell imaging of labeled H2B histones in widefield compared to light sheet excitation. The sample is observed in 3D using a 4-planes MFM setup. (C) The volumetric reconstruction of the dynamic data. (D) Histograms of the 3D apparent diffusion coefficients of 3 different nuclear factors with different dynamic behavior: DNA-bound H2B histones, Cohesin with dynamics ranging between freely diffusing and bound molecules, and CTCF that is mostly found in a bound state. (E) 3D Super-resolution imaging of Lamin-B1 in U2OS cells that are 20um above the coverslip. (F) characterizing the 3D localization precision. (G) Histogram of localization count along the cylinder show in (E) illustrating the resolution improvement in V-LSFM.

### Reaching SM regime in high density structures

Single-molecule imaging in dense structures relies on different strategies to reach single-molecule regime. The main purpose is to obtain a high signal-to-noise ratio (SNR) for the precise localization of single-molecules. Illuminating the sample with considerable light doses bleaches most of the dyes to reach a sparse distribution of emitters per frame. In dense labeling, the bleaching step helps in reducing the background noise from out-of-focus molecules that pollute the signal. To illustrate the advantage of our V-LSFM we performed instantaneous 3D imaging of single H2B histones in the nucleus of living U2OS cells, up to 20 *μm* above the coverslip surface. H2B molecules were labeled using HaloTag technology with Alexa Fluor AF647 dyes at 5nM concentration. Imaging individual histones in the nucleus of the same cell with nonspecific widefield excitation requires long exposure of the sample to high intensity illumination in order to bleach most of the dyes (Figure 2 B). By contrast, V-LSFM reduces the out of focus background and shortens the illumination time prior to SM imaging as the bleaching and imaging are performed selectively on the desired portion of the sample. Despite the high density labeling, and thanks to the volumetric selective excitation, we were able to reach the single-molecule regime by selectively illuminating the nucleus for about ten seconds with < 2 × 10^−2^ kW/cm^2^ on-sample intensity (Figure 2 C, Supplementary video 4). We followed the dynamics of single H2B histones in the nucleus, by imaging ~30 volumes of 22μm × 22μm × 2μm per second. Single-molecules were tracked in 3D for up to 900ms before bleaching. From the reconstructed trajectories, we computed the apparent diffusion coefficient *D* of H2B for trajectories that persisted more than 10 frames. As expected for H2B, the observed dynamic range matches that of molecules bound to the DNA^19,20^, showing a diffusion coefficient peak at 0.01 *μm^2^/s* (Figure 2 D).

### V-LSFM preserves the advantage of a confined excitation

To illustrate the advantage of confined excitation in 3D imaging, we imaged mouse embryonic stem cells that are considered thick and tend to grow in 3D colonies. Stem cells are sensitive to high illumination intensity dosage that is commonly used for single molecule imaging. To reach sufficient signal to noise ratio without inducing phototoxicity, single-molecule studies in stem cells were previously performed on cell monolayers and with inclined illumination^21^. Being minimally intrusive, using V-LSFM we could image SM in these delicate samples and extend the investigation to cells organized in 3D clusters.

Mouse embryonic stem (mES) cells were seeded into the soSPIM chambers. We followed the dynamics of two nuclear factors, CTCF and Cohesin, that are involved in chromatin organization. CTCF and Cohesin were labeled with JF549 using Halo-Tag technology at concentrations of 0.5 to 2nM, and trajectories were recorded for individual proteins diffusing in the nucleus (Supplementary video 5). Nuclei of different cells of the same cluster were imaged up to 30*μm* above the coverslip surface. Volumes of 22 μm × 22 μm × 2 μm were acquired in V-LSFM at an acquisition rate of ~50 volumes per second. The same nucleus was imaged over a relatively long time period, maintaining the single-molecule regime over >2 minutes. For both nuclear factors, trajectories extending up to ~700 ms were recovered. The specific physiological role of each protein was translated into different observable dynamics that were quantified by calculating the apparent diffusion coefficient distributions for trajectories longer than 10 frames. CTCF is a DNA-binding protein resulting in slow diffusivity (0.01 μm^2^/s < D < 0.1 μm^2^/s), while Cohesin presents a wider dynamic range with a fast diffusing population (*D* > 1 μm^2^/s). The observed dynamics in 3D were in good agreement with the values that were reported in previous studies performed on monolayers of cells^21^. We confirmed that the observed 2D dynamics in STEM cells are conserved in cells organized in 3D. V-LSFM provides thus a tool for extending the studies of SM dynamics into 3D in cells growing in multiple layers or colonies and avoids tracking artifacts or biases due to multiple appearances of fast diffusing molecules. (Figure 2 D).

### V-LSFM allows volumetric super-resolution imaging in dense labeling conditions

We performed single-molecule localization-based super-resolution imaging in clusters of U2OS cells that were seeded in the imaging chambers and subsequently fixed. The nuclear membrane protein Lamin B1 was imaged using the DNA-PAINT technique^22^. Individual Cy3B dyes binding events are observed with a dye concentration of 2.5 nM in the imager solution. Such value is considered high compared to typical <1nM imager concentrations used in DNA-PAINT super-resolution imaging^22,23^. 25,000 frames were acquired at a frame rate of 200ms/frame (5 volumes per second), covering a volume of 22 μm × 22 μm × 1 μm. During this time 96,000 to 188,000 localizations were obtained (Figure 2 E). To cover a larger imaging volume, the interspace between the planes was changed and the thickness of the homogeneous light-sheet was adapted accordingly to reconstruct images of 22 μm × 22 μm × 2 μm volumes with 11,0000 localizations. The total acquisition time was <1h30. The time required to image the same volumes at comparable final image quality by a sequential acquisition of 4 single plane light-sheet with a spacing of 400nm is estimated to 6h. By analyzing the multiple appearance of the same molecule on consecutive frames, we quantified the empirical localization precision to be 20 *nm* × 50 *nm* (xy × z) (Figure 2 F). Such localization precisions are comparable to those obtained with other optical sectioning methods for single-molecule imaging^5^. With such localization precision, we were able to observe details of the structure of the nuclear membrane below the diffraction limit (Figure 2 G).

## Conclusion and Discussion

The novel and completely modular V-LSFM offers a selective and accurate instantaneous volumetric single molecule imaging and tracking within extended volume far from the coverslip surface that can be adapted to various biological systems. Our approach relies on a confined and uniform intensity distribution over the thickness of the excitation light-sheet that matches the observed volume. The uniform illumination is obtained by engineering the excitation beam to form a double arc shape. Our first implementation relied on a DMD that allows a modular and fast switching between different sheet thicknesses. The diffractive nature of the DMD pixels create several challenges related to wavelength dependency of the output pattern, and, notably, a compromise between the diffraction efficiency of the different excitation wavelengths was necessary (Supplementary Information). Specifically, for a fixed light-sheet thickness, a simple opacity mask reduces the complexity of the setup and reduces light losses.

In our simulations, the double arc shape of the beam was found to yield an optimum intensity distribution along the light-sheet. However, we don’t disregard that other optimization processes might yield other possible solutions to the same problem.

Our implementation relies on a soSPIM architecture for producing the light-sheet. However, our approach can be adapted to other strategies for SPIM such as using two objectives or a tilted excitation combined with remote imaging at the expense of an anisotropic lateral resolutions.

In V-LSFM, MFM is a key element for the instantaneous volumetric imaging of the whole extent of the excited sample. In addition, MFM does not introduce any image distortion, and thus is compatible with structural widefield imaging of cells. The focus of this paper was on single-molecule imaging in single cells using confined excitation, still other approaches can benefit from the V-LSFM excitation and emission configuration for improving the resolution^24,25^. For single molecule imaging, other methods relying on PSF engineering^26,27^ for 3D SM localization can be implemented. The modular and uniform excitation we have demonstrated here can be matched to the imaging range of different PSF shapes and ensures a uniform localization precision over the covered volume.

By reducing the photobleaching in volumetric samples thanks to the confined excitation, V-LSFM is compatible with 3D tracking of individual biomolecules at relatively long-time scale. The ability to follow single molecules over large depth improves trajectory identification and avoids overestimating the proportion of particles with high diffusion coefficients due to multiple re-appearance in the depth of field of single plane imaging techniques. The selective excitation improved the DNA-PAINT super-resolution imaging in 3D. By reducing the background of freely diffusing molecules, the imager concentration was doubled compared to typical single-plane widefield acquisitions. Moreover, in contrast to 4 sequences of sequential acquisitions of 4 different planes with single-plane imaging approaches, the V-LSFM captures 4 focal planes simultaneously and thus reduces the acquisition time of 3D SMLM. Finally, V-LSFM can be applied to other super-resolution technique based on single-molecule localization^28–31^.

The results presented here demonstrate that V-LSFM offers an opportunity to perform new exciting biological investigations at the molecular scale in multi-layer systems such as cell colonies and organoids away from the coverslip.

## Methods

### Setup

The microscope setup is based on the combination of five independent modules or units, added around a Nikon Ti eclipse inverted microscope (Figure 1 A). A laser combiner (L6Cc, wavelengths: *λ*_1_ = 405 nm, *λ*_2_ = 488 nm, *Λ*_3_ = 561 nm, *λ*_4_ = 640 nm, by Oxxius) is coupled with an optical fiber of *NA_fiber_* = 0.12. At the output of the fiber, the laser beam is collimated by the lens L1 (F950FC-A, by Thorlabs, focal length *f*_1_ = 10 mm). All the modules are designed and optimized for multiwavelength imaging. The sample illumination and fluorescence collection are both performed by a single 60X, 1.27 NA water-immersion objective (Nikon CFI SR Plan Apo IR 60XC WI).

#### First module – Automated beam expander

An automated tunable beam expander, positioned after the laser combiner, regulates the size of the initial excitation Gaussian beam. The module is a compact, three-lenses compound system. The first lens is a tunable electrical lens EL1 (EL-16-40-TC-VIS-20D, optical power range −10 dpt to 10 dpt, controlled by a lens driver 4i, by Optotune). The second lens L2 (AC254-050-A, by Thorlabs), with fixed focal length *f*_2_ = 50 mm, is placed on a translation stage (LNR502/M, TravelMax by Thorlabs), controlled by a stepper-motor (BSC201, by Thorlabs). The third lens L3 (AC254-050-A, by Thorlabs) has fixed focal length *f*_3_ = 50 mm with the role of collimating the output beam. In the initial configuration *f*_*EL*1_ → ∞, the distances between the lenses are specifically 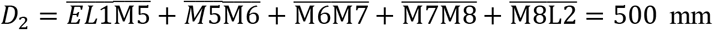 and 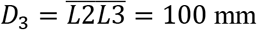. In this configuration the beam expander introduces a total magnification of *M_tel_* = 1 (Supplementary Figure 2). The design of our module imposes that at every displacement *s* of the translator the spaces between the lenses become reciprocally *d*_2_ = *D*_2_ + 3*s* and *d*_3_ = *D*_3_ − *s*. By design, the beam outcoming the beam expander is collimated when [(*f*_*EL*1_ + *f*_2_ − *d*_2_)](*d*_3_ − *f*_3_) − *f*_2_ (*f*_*EL*1_ − *d*_2_) = 0. The total magnification introduced by the module can be tuned by regulating *f*_*EL*1_ and the displacement *s*. The real magnification was calibrated by measuring the output beam diameter compared to the incoming one (Supplementary Figure 2 b-c). The excitation beam size can be regulated between 1 *μm* and 4 μm. The beam expander module is completely automatized with a LabVIEW program that integrates the drivers of the translator stage (APT software, by Thorlabs) and the tunable lens (Lens Driver Controller, by Optotune). The interface allows to introduce the desired excitation beam size, while the program automatically operates the displacement of the translator stage and the regulation of the optical focusing power of *EL*1. The automated telescope optimizes also the size of the laser beam before entering the second module.

#### Second module – Beam shaping

In our method, the excitation is confined to a selected volume of the sample to produce instantaneous 3D imaging. Therefore, it is crucial to produce homogeneous illumination over the volume of interest in order to set identical excitation and detection conditions. We designed and implemented a beam shaping module based on a DMD (DLPLCR4500EVM, by Texas Instruments) in order to produce homogenous excitation over an extended volume. The DMD is installed on a plane conjugated to the sample plane using the lens L4 (AC254-400-A, by Thorlabs) with a focal length *f*_4_ = 400 mm. The collimated excitation beam is reflected by the DMD and the shaped illumination beam is re-collimated by the microscope objective. The DMD pattern is reproduced at the focal plane of the objective.

For coherent light, the DMD acts as a blazed reflective grating, therefore diffraction and interference affect the intensity distribution after the reflection of the beam. The beam shaping of a coherent light at different wavelengths is a non-trivial problem since both these effects are wavelength-dependent. We implemented the second module with multiwavelength application in mind. We used a total internal reflection prism (TIR) and dichroic beam splitters: D1=D2 (ZT405rdc UF2, by Chroma) and D3=D4 (ZT488/640dcrb UF2, by Chroma). This configuration decouples the incident angles of the different excitation wavelengths and aligns the reflected shaped beams on the same optical path, while preserving a high-intensity despite an off-blazed condition (Supplementary Figure 3a).

In order to derive the optimal beam shaping pattern, we developed a MATLAB code that simulates the intensity of a light-sheet produced by scanning an intensity-shaped beam at the sample plane. We optimized the pattern to be produced by the DMD mirrors in order to obtain a light-sheet that could excite an extended sample volume with homogeneous intensity. The code simulates the DMD as a matrix in which each element represents a miniaturized mirror and can be switched between an ON and OFF state. The original Gaussian beam and the DMD pattern are superimposed by matrices multiplication. The simulation considers the diffraction limit imposed by the physical size of the imaging optics. By exploiting the radial symmetry of the intensity of a Gaussian beam, we found that a DMD pattern shaped as a double arch with a dark elliptical central area produces a homogeneous light sheet when teetered (Supplementary Figure 5,6).

Taking into consideration the physical size of the DMD micromirrors, we found the correspondence between the parameters of the pattern and the excitation volume produced by scanning the shaped beam (Supplementary Figure 7). By scanning the double-arch modulated beam to excite a uniform fluorescent layer, we demonstrated that we can produce an excitation volume with homogeneity intensity >80% in the range of 2 to 5 μm (Supplementary Figure 8-10).

#### T*hird module – Beam steering*

The third module is dedicated to steering the beam and controlling the light-sheet position and parameters at the sample plane. This module is based on the soSPIM beam steering unit described in ref ^13^ where three relay systems composed by achromatic lenses originate three planes that are optically conjugated to the back focal plane (BFP) of the objective. The systems are composed of lenses L5 and L6, L7 and L8 (*f*_5_ = *f*_6_ = *f*_7_ = *f*_8_ = 50 mm, AC254-050-A, by Thorlabs), and L9 and L10 (*f*_9_ = 150 mm, AC254-150-A, by Thorlabs, *f*_10_ = 200 mm, AC254-200-A, by Thorlabs). A tunable, electrical lens EL2 (Custom EL-10-30, by Optotune; focal lengths from −80 mm to +1000 mm) is placed on the first conjugated plane. This lens introduces a defocusing of the excitation Gaussian beam allowing to adjust and maintain the position of the light-sheet’s waist compared to the sample position in relation to the mirror. Two galvo-mirrors *GM_x_* and *GM_y_* (ScannerMAX 506 actuators, by Pangolin, with dielectric mirrors, by Chroma) are placed respectively on the second and third planes conjugated to the objective’s BFP. In this way, the mirrors control the tip-tilt angle of the excitation beam at the back focal plane of the objective, and thus regulate the position of the light-sheet at the sample plane. After reflection onto a 45° mirror oriented along the y-axis in the sample plane, *GM_X_* is responsible of the *z* positioning of the light-sheet, while *GM_y_* produces the fast steering of the beam in the y direction. In practice, the angle of the light-sheet scanning direction can be adjusted using the two galvanometric mirrors releasing the alignment constraints of the device onto the microscope stage. Finally, a mirror M13, placed at a conjugated plane to the sample plane, controls the tip-tilt of the excitation beam at the sample plane. This is equivalent to displacement of the beam in the BFP of the objective and allows to compensate for 45° mirror tilt in order to maintain a light-sheet perpendicular to the optical axis of the objective. The M13 mirror is placed at equal distance *f*_7_ = *f*_8_ = 50 mm between L7 and L8.

#### Fourth module – Microfabricated sample holder for soSPIM imaging

The fabrication protocol of the soSPIM devices used in this study is described in detail in Reference ^13^. Briefly, a silicon mask is first etched using a combination of anisotropic wet etching and deep-reactive ions etching processes to create respectively 45° slanted surfaces and microwells of various dimensions (24×24 μm^2^, 40×40 μm^2^, 60×300 μm^2^) along those surfaces. The silicon mask is then replicated in a secondary PDMS mold (Sylgard 184, Dow Corning), which in turn is used to reproduce the silicon mask features onto cleaned #1.5h coverslips by capillary filling of a UV-curable and index match polymer (BIO-134, MyPolymer). At this stage, the index-matched polymer could be mixed with 100 nm nano-diamonds (NDNV100 nmMd10ml, Adamas Nanotechnologies INC.) suspended in acetone to provide fiduciary markers homogenously spread in the volume all around the well. The nano-diamond concentration was experimentally adjusted to ensure the presence of at least one fiducial maker within any 4 μm deep volume in the vicinity of each well (within a radius of ≈ 50 μm). After polymer curing with the device fully immersed in water (20 min using a 500-W arc lamp, 66902, Newport), the PDMS mold is peeled-off and the coverslip coated with a thin layer of gold by plasma sputtering in a vacuum chamber (JFC-1600 Auto Fine Coater, JEOL) to make the 45° surfaces reflective. The mirrored surfaces are then protected with a thin layer of UV-curable polymer (NOA 73, Norland Product) by capillary filling while the microwells are protected by a flat PDMS piece followed by 2 min UV insolation. Finally, the flat PDMS piece is peeled off, and the unprotected metal coating is removed by wet etching (gold etchant, 651818, Sigma) leaving the slanted surfaces coated with gold and the microwell uncoated for optimal light transmission. Finally, before use, the devices are immersed in pure ethanol (Sigma) and insolated with UV for 2h to reduce the polymer auto-fluorescence and ensure minimal background light.

#### Fifth module – Multifocus microscopy

Finally, the MFM module^16,17^, essentially composed of three custom-designed optical elements, was installed on the detection path. The tube lens of the microscope TL (*f_TL_* = 200 mm) and an achromatic relay lens L11 (*f*_11_ = 150 mm, AC254-150-A, by Thorlabs) produce a secondary pupil plane, or Fourier plane, where a multifocus grating (MFG) is placed. The MFG (custom-made by Creative Microsystems,) performs two distinct functions. First, it splits the fluorescence signal into separate paths, each one associated with a diffraction order (*m_x_*, *m_y_*). The central grating motif of the MFG maximizes the diffraction efficiency and ensures a uniform distribution of the fluorescence light in the desired diffraction orders^18^. Second, it adds a specific defocusing power to each diffraction order. A combination of multifaceted prism and blazed grating correct any chromatic aberration created by the MFG. A final lens L12 (*f*_12_ = 200 mm, ACT508-200-A, by Thorlabs) refocuses the several diffraction orders on different parts of the imaging camera (iXon Ultra897, 512×512 active pixels, physical pixel size 16 μm × 16 μm, by Andor). As such, an array of images of various focal planes with constant focal step Δz are tiled side by side and imaged instantaneously on the same camera.

### Multifocus Grating design

A multifocus grating (MFG) is at the core of multifocus microscopy. The MFG is a binary phase grating etched into a glass substrate that is conceived for a specific objective and enables a predefined plane spacing. The repeating motif is optimized in order to yield a high diffraction efficiency into the desired diffraction orders while preserving a uniform light distribution. First implementation of MFM relied on a 9 plane MFG configuration and was used with a Nikon 1.4NA oil immersion objective. Here we have designed a new set of gratings conceived to work with a Nikon 1.27NA 60x water immersion objective and yielding several plane separations (250nm and 500nm) with dual color properties^18^. The motif in these gratings were adjusted to produce 9 or 4 planes. The simulation of the central motif and the distortions were performed using a custom-made code. The gratings were produced by photolithography based on the provided designs (SILIOS Technologies, France).

### The soSPIM imaging software

We slightly modified the software from ref ^13^ that controls the laser beam steering unit. It runs using MetaMorph acquisition environment (Molecular Devices) that controls the microscope component (XY and Z stages, microscope filters turret) and handles the 3D acquisition process. The soSPIM software allows synchronizing the light-sheet displacements with the acquisition process to ensure optimal 3D imaging over the whole depth of the imaging device. Briefly, after defining the 45° mirror axis, a two-step calibration process enables to synchronize the displacement of the light-sheet position with the waist location and the objective axial position to ensure that the excitation and detection planes remain permanently superposed. The software allows also to adjust the light-sheet length and orientation. For our method, we specifically implemented a dedicated z-scanning module along the mirrors depth to allow for axially extended excitation volume that match the MFM volume detection.

### Adjustment of the DMD pattern

Due to a limited mechanical precision and the presence of multiple optical elements inserted between the DMD and the observation camera, the frames of the DMD and of the camera were slightly misaligned. As the teetering of the excitation beam is commonly performed along the dimensions of the camera frame, it was necessary to compensate for any rotation. Using the uniform fluorescent layer, a pattern of regular grid of bright points was applied on the DMD. The excitation grid was imaged on the camera and found to exhibit a slight tilt compared to the camera frame. A home-made MATLAB codes identified the center of the peaks and found the appropriate transformation to be applied on the DMD pattern in order to match the main axis of the camera frame. This transformation was applied on any pattern that was applied to the DMD, including the double arc pattern, in order to match the camera main frame (Supplementary Figure 7).

### Uniform fluorescent layer

To evaluate the illumination uniformity and size produced by modulating the excitation light-sheet or by steering a Gaussian beam in two dimensions, we fabricated a homogenous fluorescent layer. We mixed a fluorescent dye in a 5% polyvinyl alcohol (PVA) solution. A 30 μl drop of the fluorescent solution was placed in the middle of a coverslip, and spin-coated at 2000 rpm for 30s producing a 400 nm thick layer. The coverslip was cured in an oven at 80 degrees for 2 hours.

### Cell culture and labeling

For imaging the 3D dynamics of H2B histones, we used human osteosarcoma (U2OS) cell line. For the super-resolution experiments, we used the U2OS cell line with a stable insertion of EGFP on NUP96 nuclear protein (U2OS-NUP96EGFP) purchased from CLS cell line services.

The cell culture protocol is the same for U2OS and U2OS-NUP96EGFP. The cells were grown in DMEM (11880, by Thermo Fisher Scientific) + 1% Glutamax + 1% Penicillin-Streptavidin supplemented with 10% fetal bovine serum (26140079, by Thermo Fisher Scientific). Cells were maintained at a density of 0.2 — 1.5 × 10^5^ cells/cm^2^ at a temperature of 37° in the presence of 5% CO2. Cells passed regular tests for ruling out any mycoplasma infection.

For the observation of the 3D dynamics of CTCF and Cohesin we used mouse embryonic stem (mES) cells. The cells lines used in this study were genetically modified to stably express CTCF and Cohesin carrying a Halo-Tag. Cells were cultured in DMEM+Glutamax (10566-016, by Thermo Fisher Scientific) supplemented with 15% fetal bovine serum (S1810-050, Lot S15642S1810, by DUTSCHER), 550mM b-mercaptoethanol (21985-023, by Thermo Fisher Scientific), 1mM sodium pyruvate (11360-070, by Thermo Fisher Scientific) and 104U of Leukemia Inhibitory Factor (ESG1107, by Millipore). Cells were maintained at a density of 0.2-1.5 10^5^ cells/cm^2^ by passaging using TrypLE (12563011, by Thermo Fisher Scientific) every 24h - 48h. Cells were grown on 0.1% gelatin-coated dishes (ES-006-B, by Millipore) and incubated at a temperature of 37° in the presence of 5% CO2. The culture medium was changed daily. Cells passed regular tests for ruling out any mycoplasma infection. Cells were seeded at a density of 3· 10^5^/cm^2^ the day before the experiments in Fluorobrite DMEM (A1896701, by Life Technologies SAS-Thermo Fisher Scientific).

To observe the dynamics of H2B histones, confluent U2OS cells seeded in a Petri dish were transfected using X-tremeGENE HP DNA transfection reagent (6366236001, by SigmaAldrich) to express H2B protein carrying a Halo-tag. Labeling was performed after 24h of transfection by incubating cells for 15 minutes in 1 mL solution of Halo-conjugated AlexaFluor 647 dyes at a concentration of 5 nM in enriched DMEM. Cells were then washed with enriched DMEM few times and transferred to the soSPIM chip.

To observe the dynamics of CTCF and Cohesin in stem cells, confluent mES cells were cultured in a Petri dish with Fluorobrite DMEM. The cells were labeled with Halo-conjugated JF549 by 15 minutes incubation in 1 mL solution of the dye at different concentration in Fluorobrite DMEM. After washing with Fluorobrite DMEM, the cells were seeded in a soSPIM chip.

### DNA-PAINT sample preparation

Cells were transferred from the culturing flask into a soSPIM chip, in which previously enriched DMEM was incubated for 30 minutes under vacuum. After 15 minutes, cells were precipitated in the wells and could be fixed. For the fixation, cells were incubated in a solution of 4% PFA in PBS for 10 minutes at room temperature. After washing with PBS, cells were incubated for 10 minutes in a solution of PBS + 150 mM glycin (26-128-6405-C, by EUROMEDEX) to quench the autofluorescence of the PFA. After another washing, cells were permeabilized for 10 minutes in a solution of PBS + Triton™ X-100 at a concentration of 2% (volume/volume). Cell were washed and then incubated in the blocking buffer (3% BSA (A9418, by Sigma-Aldrich) + 0.1 mg/mL of sheared salmon sperm DNA (AM9680, by Thermo Fischer Scientific) + 1% dextran sulfate sodium (D4911, by Sigma) in PBS) for 90 minutes. The blocking buffer was removed and cells were incubated with recombinant anti-Lamin B1 primary antibody (ab229025, by abcam) diluted 1:250 (volume/volume) in the blocking buffer. Cells were washed with the blocking buffer and were incubated for 60 minutes with the secondary antibody (included in MASSIVE-SDAB 1-PLEX by Massive photonics) diluted 1:500 (volume/volume) in the blocking buffer. After washing the sample with blocking buffer and imaging buffer (included in MASSIVE-SDAB 1-PLEX by Massive photonics), the solution with the DNA imager strands were added to cells. The optimized imaging solution was prepared with imaging buffer and an imager strands that is complementary to the secondary antibody (included in MASSIVE-SDAB 1-PLEX by Massive photonics) at a concentration of 2.5 nM. After imaging, cells were stored in blocking buffer at 4°C.

### Seeding cells in the soSPIM wells

To seed pre-transfected and labeled cells in the soSPIM sample holder, first, the chip was covered with culturing medium and incubated under vacuum for 30 minutes to remove any air bubbles trapped in the wells. Cells were transferred to a 15 ml falcon using TrypLE (12563011, by Thermo Fisher Scientific) and centrifuged at 300 g for 3 minutes. Cells were resuspended in 200 μl of culture medium, then the soSPIM microfabrication was covered with a volume of 100 *μl* of cells solution that was delicately placed to form a drop on top of the chip. After 15 minutes incubation at 37°, live cells are precipitated in the cavities. The device was then be washed with culture media to remove any cells lying outside of the wells before to be used either for imaging or cell fixation and labelling before imaging.

### Image reconstruction

We followed the same process for image reconstruction as mentioned previously^17^. We recall the main principle here: multicolor beads deposited on a coverslip are imaged in MFM using widefield excitation. As the beads are displaced axially using a piezo stage, they appear consecutively in focus on the different MFM planes. The obtained stack serves as a reference to generate the transformation matrices necessary to align the different MFM planes and for reconstructing the instantaneous Zstack of each time frame.

### Data analysis and visualization

Single molecule acquisition sequences were reconstructed as Zstacks using the above-mentioned approach. As single molecule PSFs are sampled simultaneously on several focal planes, a routine based on 3D gaussian fitting allows to recover precisely the center of each molecule^17^.

Localization data was filtered using a custom-made MATLAB code for bad detections by thresholding large standard deviations or out of bounds axial positions.

To correct the drift in super-resolution acquisitions, two strategies were used. A first strategy relies on the introduction of fiducial markers^32^ in the polymer forming the soSPIM device. The particle was inserted randomly at a given concentration with the aim to ensure the presence of at least one marker within any 4 μm deep volume around each well. Drift correction is achieved off-line by 3D tracking the fiduciaries. In case no fiducial marker was present within the probed volume, a second approach based on cross correlation was used to correct the drift^33^. The 3D cross correlation routine was extracted from the SMAP software describe in ^34^.

For dynamic data in live cells, we used the Utrack algorithm to track single molecules in 3D ^35^. The obtained trajectories were later filtered to rule out tracks with less than 10 time points, and mean square displacement (MSD) was calculated for each trajectory. Apparent diffusion coefficients were computed for each trajectory by linear-fitting the 5 first points of the MSD using MSD(t) = 6Dt+b for the 3D case.

The localizations were visualized in the Genuage software in desktop mode^36^. Genuage was used to produce the supplementary videos.

## Acknowledgments

BH acknowledges funding from the Fondation pour la Recherche Médicale (FRM; DEI20151234398), the Agence National de la recherche (ANR-19-CE42-0003-01), the LabEx CelTisPhyBio (ANR-11-LABX-0038, ANR-10-IDEX-0001-02), the Institut Curie, Agence pour la Recherche sur le Cancer (ARC Foundation), DIM ELICIT and from ITMO Cancer of Aviesan on funds administered by Inserm (grant N° 20CP092-00). BH, RG and JBS recognize the support of France-BioImaging infrastructure grant ANR-10-INBS-04 (Investments for the future).

TG acknowledges funding from the European Union’s Horizon 2020 research and Innovation program under the Marie Sklodowska-Curie Grant Agreement No 666003 and the Fondation pour la Recherche Médicale under the *Fin de these 2020 program* (FDT202001010813).

RG and JBS acknowledge the soLIVE funding from the Agence National de la Recherche (ANR-16-CE11-0015-01).

We thank Luke Lavis for providing the JF dyes, and Elphege Nora for sharing the STEM cells that were imaged in this paper. We thank Romain Laine for useful discussion, and Hisham Forriere for the sharing labeling protocols and providing feedback.

We dedicate this paper to the memory of Maxime Dahan.

## Author contributions

BH conceived the idea and directed the project. BH designed the MFGs. TG and BH mounted the optical setup and acquired the data. BH, YF and TG wrote the analysis routines. TG and HRZ analyzed the data. TG prepared the samples. TG and LC prepared the STEM cells samples. RG, JBS and VV provided the soSPIM optics and sample holders. JBS programed the soSPIM control routines. TG and BH wrote the paper. All the authors participated in planning the experiments and contributed to the writing of the paper.

## Data and code availability

All data is accessible upon request

