## Supplementary Information for "Selective volumetric excitation and imaging for single molecule localization microscopy in multicellular systems"

### Multifocus Grating

The multifocus grating (MFG) is a phase mask placed conjugated to the back focal plane of the objective. It is a phase only grating that is etched into a fused silica substrate.

The main motif of the grating is designed to produced a desired distribution of the diffracted light while aiming for maximum intensity values (Efficiency) and homogeneous light distribution between the desired diffraction orders (Uniformity). The design is based on an iterative process using a custom-made MATLAB script. The output grating design is also optimized for two distinct colors as described before[1]. Here, the photolithography process was taken into consideration as possible dilation and erosions could impact the output structure and thus the diffraction efficiency. In addition, the motif was corrected manually by rounding sharp edges and widening thin structures while ensuring a correct diffraction performance.

The main designs for producing 9 and 4 planes with dual color capabilities are presented in **Supplementary Figure 1**.

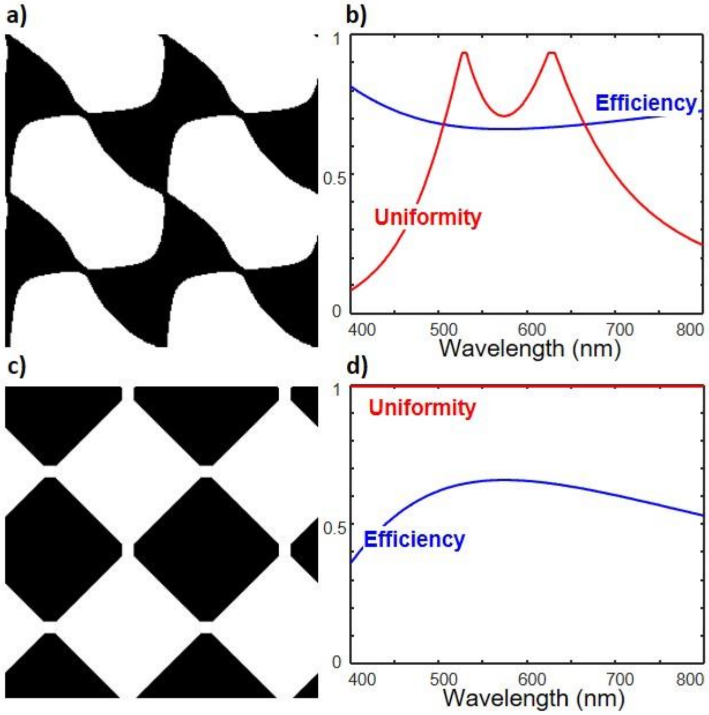


***Supplementary Figure 1 :*** *Multifocus Grating design. (a) the design of the 9 planes MFG grating that was used in this work. The motif was adjusted to avoid sharp edges. (b) shows the corresponding diffraction efficiency and uniformity as a function of the imaging wavelength. The grating was designed to work with 2 different imaging wavelengths at 525nm and 620nm. (c) the motif design of the 4 planes MFG with the corresponding spectral characteristics (d). The uniformity and Efficiency are relatively conserved over a large spectral range.*

### Automatized Beam expander

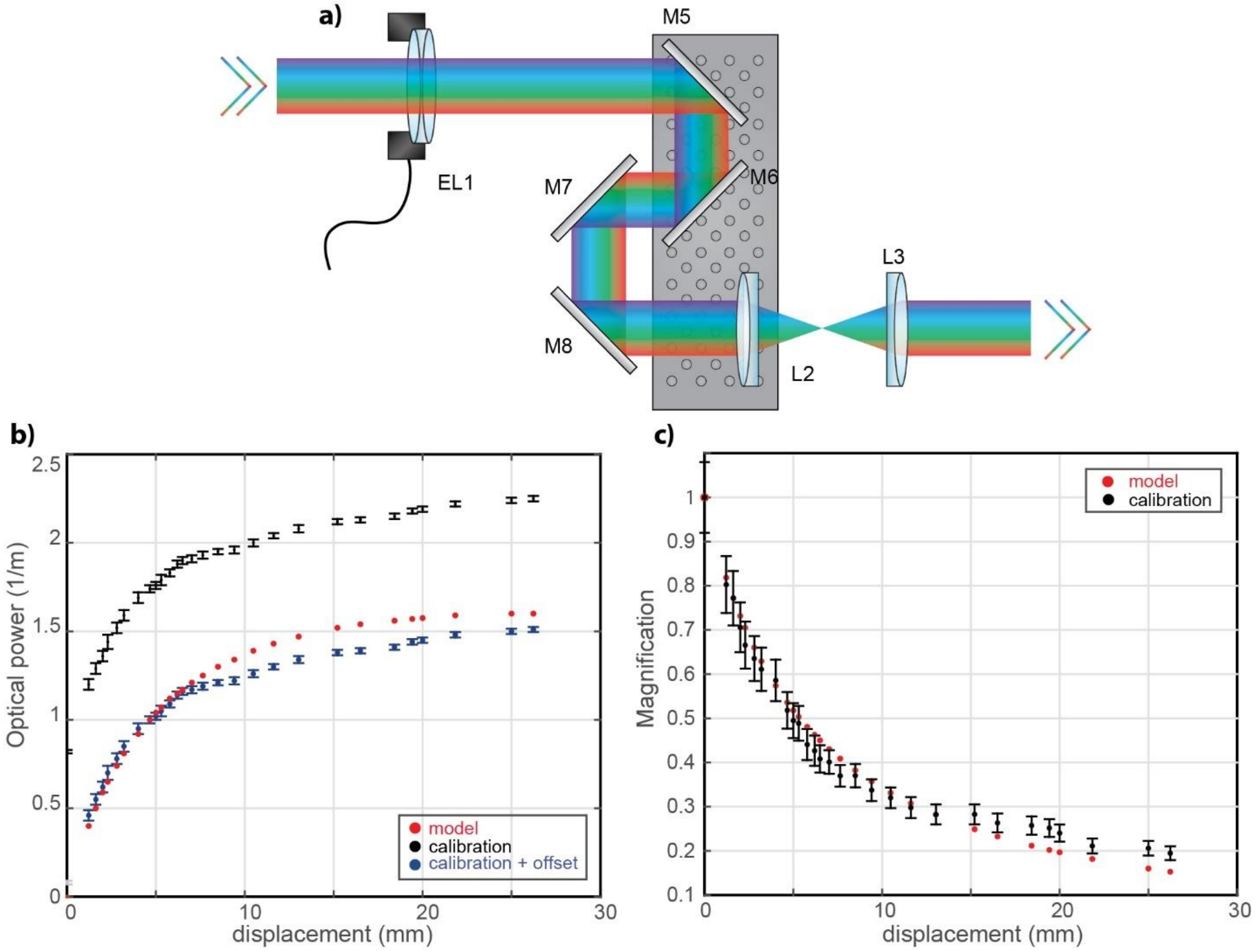


***Supplementary Figure 2 :*** *Automatized beam expander. (a) schematic presentation of the 3-lens system for producing an automatized beam expander. (b) The required optical power of EL1 as a function of the translation stage displacement comparing the theoretical calculation and the measured values. (c) The realized demagnification factors as a function of the translation stage displacement.*

The basic principle of an automatized beam expander relies on a combination of two lenses, with variable focal lengths. To output a collimated beam the distance between the two lenses have to be adjusted to match the sum of the two focal lengths.

The required demagnification in our experiments must range between 1 and 0.1. Realizing such large range of demagnifications is difficult with only two lenses with variable focal lengths. We combined an electrical lens (EL1) with a variable focal length compound lens. The later is formed by two lenses (L2 and L3) with fixed focal length. L2 is placed on a translation stage, the displacement of the translation stage affects the distances between the different lenses.

We define $f_{EL}$, $f_2$ and $f_3$ the focal lengths of $EL$, $L2$ and $L3$, respectively. We call $d_2$ the distance between $EL$ and $L2$ and $d_3$ the interspace between $L2$ and $L3$. The front focal length of the three-lenses compound system is

$$F = \frac{f_3(d_3 - \Phi)}{d_3 - (f_3 + \Phi)}$$

Where

$$\Phi = \frac{f_{EL}(d_2 - f_2)}{d_2 - (f_1 + f_2)}$$

is the back focal length of the module.

The output beam is collimated if $F \to \infty$, thus if

$$[d_2 - (f_{EL} + f_2)](d_3 - f_3) - f_{EL}(d_2 - f_2) = 0$$

The design of our module imposes that for a displacement $s$ of the translator, $d_2 = D_2 + s$ and $d_3 = D_3 - 3s$. $D_2$ and $D_3$ are the initial distances between $EL$ and $L2$ and $L2$ and $L3$ respectively (**Supplementary Figure 2Error! Reference source not found.-a**). Moreover, the translator can produce a maximum displacement of 50 mm, while the optical power range of $EL$ is -10 dpt to 10 dpt. Imposing these boundary conditions in the above equation gives the combination of displacement $s$ and focal length $f_{EL}$ for all the magnification values $M_{tel}$ that the module is required to produce. A solution was found for $f_2 = f_3 = 50$ mm (AC254-050-A, by Thorlabs) if positioned such that $D_2 = 500$ mm and $D_3 = 100$ mm.

The theoretical predictions were compared to the experimental values in (**Supplementary Figure 2-b-c**).

### Working with a DMD

#### Physical principle

A DMD is a micro-optoelectrical system composed by a matrix of electro-mechanical miniaturized mirrors, each can be independently rotated by a tilt angle ±t. The intensity profile of the reflected light is thus modulated according to the orientation of the different mirrors.

Using DMD in combination with coherent light sources originates non-trivial problems for the design of the microscope system, especially for multiwavelength applications. The DMD is to be considered as a reflective blazed grating, and diffraction effects must be taken in account for the optimal design of the setup. The DMD can be described as a two-dimensional reflective blazed grating. The theoretical basis can be found in [2–4]. Here we recall some basic principles.

An advantage of a blazing grating is to direct most of the diffraction intensity in a single diffraction order under a specific configuration. A diffraction order is in the blazing condition if it overlaps with the peak of the envelope function of the grating. When the excitation beam is incident on the DMD with an angle $\theta_i \neq 0$, the $m^{th}$ diffraction order satisfies the blazing condition if

$$m = 2\frac{d}{\lambda}[\sin(t)\cos(\theta_i - t)]$$

is an integer number, $t$ being the tilt angle of the mirrors **Supplementary Figure 3 - a**.

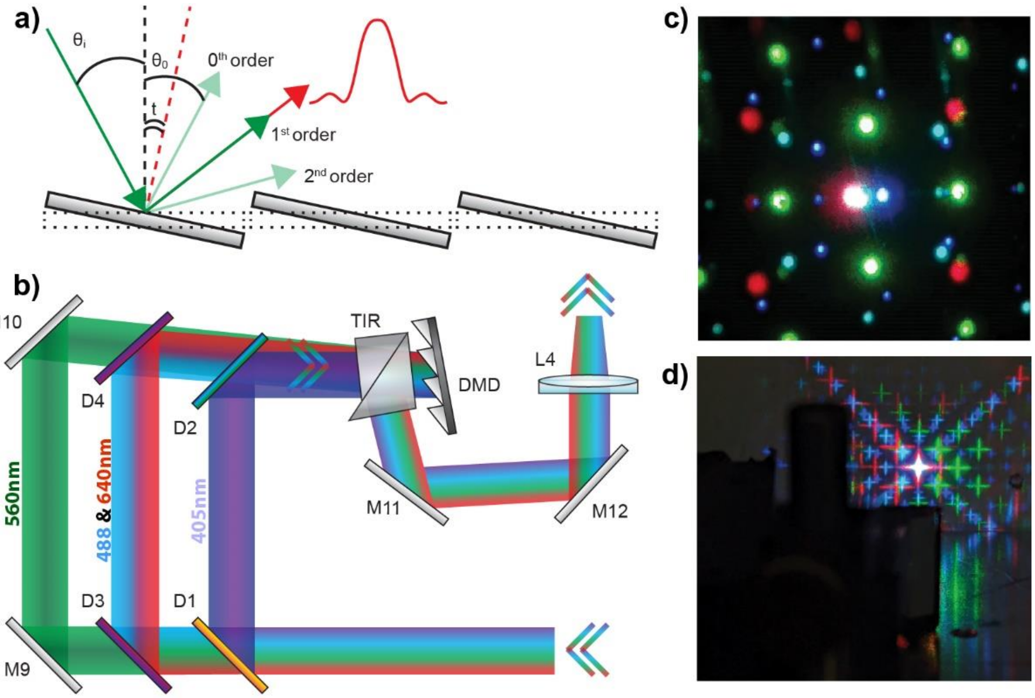


***Supplementary Figure 3 :*** *Correcting the wavelength-dependent diffraction on the DMD. (a) Schematic presentation of the tilted pixels of the DMD acting as a blazed grating and showing the different incidence and diffraction angles compared the tilting angle. The diffraction orders are wavelength dependent. (b) Compensating the blazing condition for the different excitation wavelengths by a combination of several dichroic mirrors. (c) the non-overlapping diffracted beams before correction. (d) the different overlaying excitation beam when applying (b). Here the DMD pixels were shaped as a cross to facilitate the visualization.*

In our case, $d = 10.8$ µm, $t^{\pm} = \pm 12°$ and the excitation wavelengths are $\lambda_1 = 405$ nm, $\lambda_2 = 488$ nm, $\lambda_3 = 561$ nm, $\lambda_4 = 640$ nm. For $\theta_i \sim 20°$, it is possible to determine the blazed diffraction orders for each wavelength, that are $m_1 = 11$, $m_2 = 9$, $m_3 = 8$, $m_4 = 7$, respectively. In reality, the two-dimensional diamond geometry of the grid and the rotational axis built on the diagonal of each miniaturized mirror results in a different, more complex, prediction of the DMD blazing conditions. In fact, we observed that if all the wavelengths $\lambda_1$, $\lambda_2$, $\lambda_3$ and $\lambda_4$ are incident on the DMD with the same incident angle $\theta_i$, the respective refractive angles $\theta_{m1}, \theta_{m2}, \theta_{m3}$ and $\theta_{m4}$ are different. Moreover, we observed that, at a given $\theta_i$ for which $\lambda_1, \lambda_2$ and $\lambda_4$ satisfy the blazing condition, $\lambda_3$ is off-blazed. Specifically, for $\theta_i \sim 20°$, the observed blazed order for $\lambda_1, \lambda_2$ and $\lambda_4$ are $m_1^{ob} = (10, 10)$, $m_2^{ob} = (8,8)$, $m_4^{ob} = (6,6)$, while for $\lambda_3$, the light is mostly distributed in the orders (6,6), (6,7), (7,6) and (8,8). Therefore, for the design and optimization of the beam shaping module we had to face two issues. First, we wanted the shaped beams at different wavelengths to be refracted with the same angle, so that they would propagate on the same optical path. Second, we wanted to optimize the energy distribution in the diffraction orders to excite the sample with high illumination intensity for performing efficient single-molecule imaging.

### Correcting the wavelength-dependent diffraction

In order to meet the two mentioned requirements, we found a compromise between the blazing conditions and the efficiency of diffraction for the different excitation wavelengths. Specific dichroic mirrors separate the optical path of $\lambda_1$, $\lambda_2$, $\lambda_3$ and $\lambda_4$ before the DMD. In this way it is possible to change specifically the incident angle of each wavelength and the optical paths of the diffracted beams overlap. In addition, a total internal reflection (TIR) prism was introduced to reduce the number of required dichroics. The TIR prism was included in the original DLP LightCrafter 4500 spatial light modulator. When placed correctly in front of the DMD grid, the effect of the TIR-prism is to overlap the diffracted orders containing the maximum energy of the wavelengths $\lambda_2$ and $\lambda_4$. The same optical path was thus possible for $\lambda_2$ and $\lambda_4$. In practice the dichroic beamsplitter $D1$ (ZT405rdc UF2, by Chroma) reflects $\lambda_1$ and transmits $\lambda_2, \lambda_3$ and $\lambda_4$. The multiband dichroic beamsplitter $D3$ (ZT488/640dcrb UF2, by Chroma) reflects $\lambda_2$ and $\lambda_4$ and transmits $\lambda_3$. In this way, the incident angle $\theta_i^1$ and position of $\lambda_1$ in respect to the DMD grid can be regulated by $D1$ and a second identical beamsplitter $D2$. In the same way, the incidence of $\lambda_2$ and $\lambda_4$ can be regulated by $D3$ and a second equal multiband dichroic beamsplitter $D4$. The regulation of the incidence of $\lambda_3$ is done using two mirrors $M9$ and $M10$ **Supplementary Figure 3 b-c**.

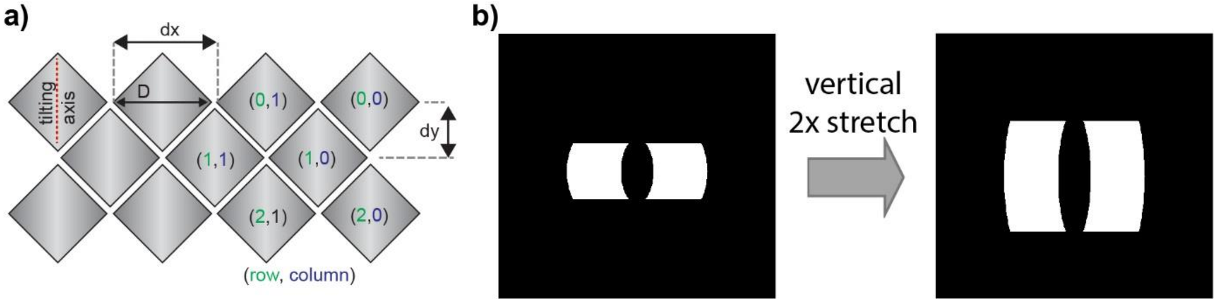


***Supplementary Figure 4:*** *Correcting the effect of the diamond arrangement of the DMD-pixels. (a) the pixel assignment of the DMD results in a* $2\times$*compression of the applied pattern along the vertical direction. (b) To compensate this effect the desired pattern is stretched two folds along the vertical dimension before application on the DMD.*

### Double arc beam

#### Simulation, parameters and propagation

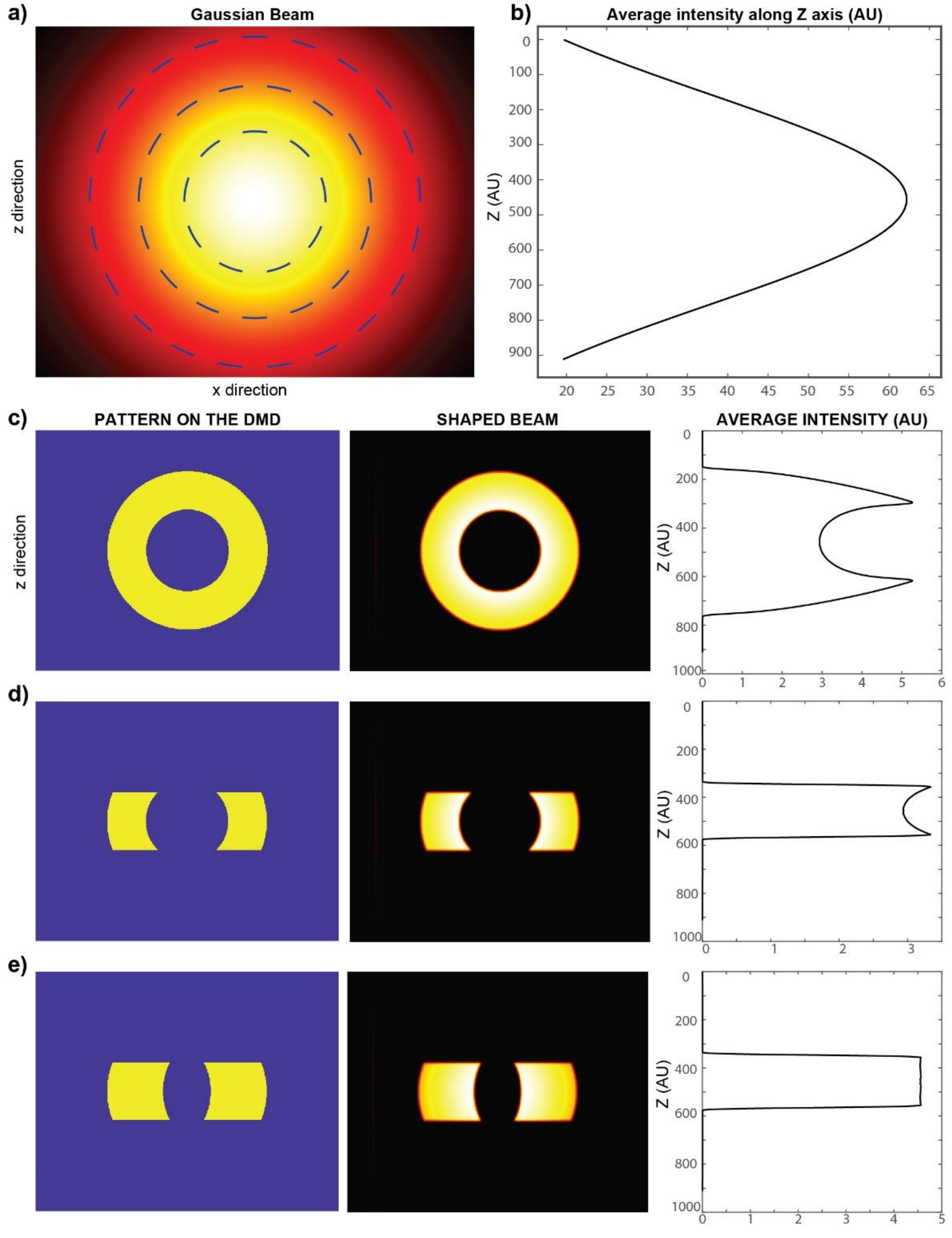


***Supplementary Figure 5 :*** *Designing the uniform axial excitation. (a) The Gaussian intensity distribution of the excitation beam. Dashed lines present the zones with equal intensities that are radially distributed around the center of the beam. A light sheet produced by scanning a Gaussian beam results in a Gaussian intensity profile along the z axis (b). The rational of finding a suitable beam profile is illustrate in (c-e). First column presents the physical ON-OFF mask to be applied on the beam. The middle column shows the resulting excitation beam after the mask at the objective focal plane and taking into consideration the diffraction of light. In the right column, the axial intensity profile for the corresponding scanned beams are plotted. The idea relies on the radial symmetry of the intensity around the beam center. To realize a uniform axial excitation profile, it is necessary to remove the central intensity peak of the beam. A circular mask of the center reduces the non-homogeneity of the excitation profile (c). An elliptical central mask reaches the desired uniformity (e).*

The intensity profile of a laser beam follows a Gaussian profile that can be described by

$$I(x,z) = I_0 e^{-\frac{(x^2+z^2)}{2\sigma^2}}$$

$(x, z)$ refer to the coordinates within the frame perpendicular to $y$, the laser beam propagation direction. The intensity profile $I$ spreads over a standard deviation of $\sigma$, and $I_0$ represents the peak intensity of the beam at the center (0,0).

The Gaussian profile is radially distributed and the intensity profile can be rewritten in the polar coordinates as

$$I(r) = I_0 e^{-\frac{(r^2)}{2\sigma^2}}$$

$r$ is the radial coordinate and is equal to $r = \sqrt{x^2 + z^2}$.

A typical light sheet is formed by scanning the Gaussian beam laterally along a linear direction (along the $x$ direction). The obtained axial intensity profile can be estimated by integrating the 2D gaussian profile along discreate lines parallel to the beam scanning direction. Along a given axial location $z_0$, the intensity profile of the Gaussian beam is given by $I(x,z_0) = I_0 e^{-\frac{(x^2+z_0^2)}{2\sigma^2}}$ which integral represents the amount of intensity of the light sheet as seen by a molecule at $z_0$ position from the center. This integral can be expressed as

$$I_{Tot}(z_0) = I_0 \sigma\sqrt{2\pi} e^{-\frac{z_0^2}{2\sigma^2}}$$

, which shows a Gaussian intensity profile as a function of $z_0$ **Supplementary Figure 5 a-b**.

In order to transform the beam profile into a uniform distribution our idea relies on the symmetry of a Gaussian beam. The beam is radially distributed in concentric rings of equal intensities. As a first tentative, it could be sufficient to keep an equal number of rings per axial position over the axial range of interest to obtain an equal distribution of the excitation intensity along the $z$ axis. This is accomplished by clipping the Gaussian intensity profile by (i) an inner disk of $r_1$ radius, (ii) an outer diaphragm of $r_2$ radius ($r_1 < r_2$) **Supplementary Figure 5** c, and (iii) a rectangle of $z'$ height **Supplementary Figure 5 d**. The intensity of the remaining pattern along a given z plane can be written as

$$I'(z) \propto e^{-\frac{z^2}{2\sigma^2}} \times \left[Erf\left(\frac{\sqrt{r_2^2 - z^2}}{\sigma\sqrt{2}}\right) - Erf\left(\frac{\sqrt{r_1^2 - z^2}}{\sigma\sqrt{2}}\right)\right]$$

The corresponding intensity profile (**Supplementary Figure 5 d**) while reduces the intensity difference along z still exhibits sharp peaks at the edge of the light sheet mainly due to a variation in the width of the pattern along the z axis (variables of the *Erf* function).

As a second strategy, we decided to change the ellipticity of the clipping mask instead of a perfect circular symmetric one. We developed an algorithm for optimizing the ellipticity of the clipping masks in order to obtain the desired uniform distribution. An optimum optimization result is shown in **Supplementary Figure 5 e**. The algorithm considers the diffraction of light due to the optics when reaching the sample plane. The simulated intensity mask results in a >90% uniformity of the light sheet over the axial range of interest. For different light sheet thicknesses, new masks were simulated resulting in different ellipticities optimized for a specific Gaussian beam size.

The analytical expression of the intensity distribution in this case can be expressed by

$$I'(z) \propto e^{-\frac{z^2}{2\sigma^2}} \times \left[Erf\left(\frac{x_2(z)}{\sigma\sqrt{2}}\right) - Erf\left(\frac{x_1(z)}{\sigma\sqrt{2}}\right)\right]$$

where $x_1$ and $x_2$ refer to the lateral limits of the inner and outer ellipses defined as function of z shown in **Supplementary Figure 6**.

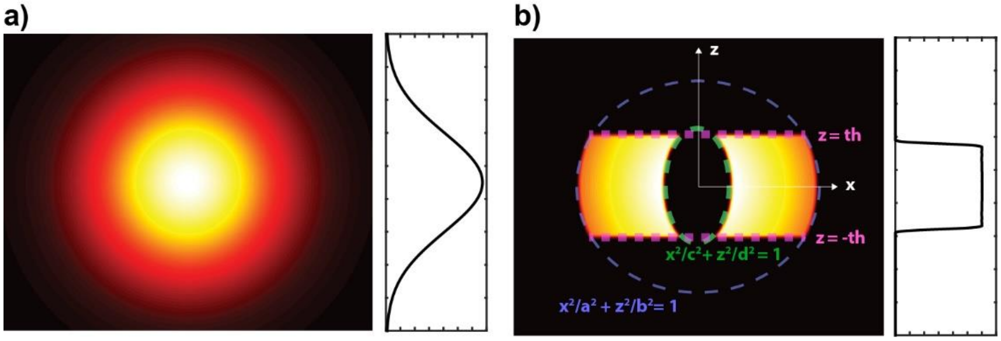


***Supplementary Figure 6 :*** *Designing the excitation beam. The Gaussian beam (a) is masked by three masks. (b) First, the beam is clipped by a linear slit of dimension* $2 \times th$ *along the z axis. This slit defines the axial extent of the excitation beam. Second, an elliptical mask blocks the beam outside the region defined by* $x^2/a^2 + z^2/b^2 = 1$ *. Third, an elliptic mask blocks the beam inside the region defined by* $x^2/c^2 + z^2/d^2 = 1$*. The parameters of the 3 masks* $(th, a, b, c, d)$ *are found by simulation for a specific axial beam size and to yield a desired light sheet thickness.*

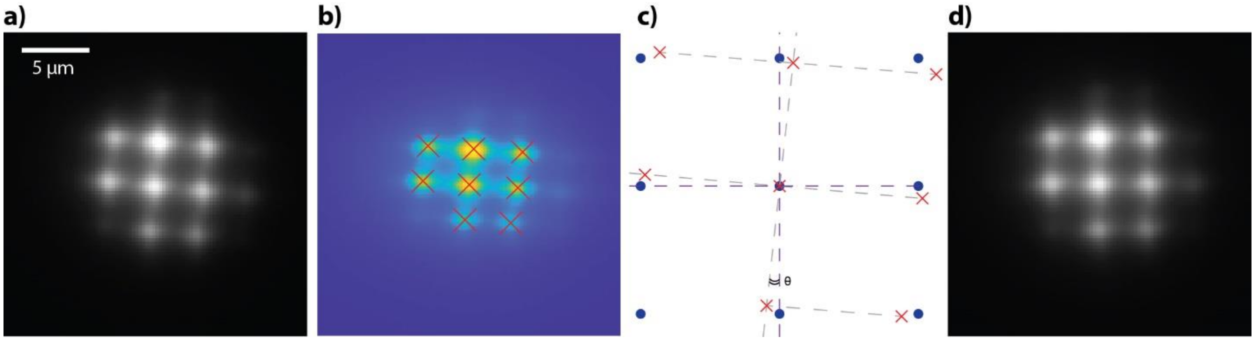


***Supplementary Figure 7 :*** *Aligning the DMD and the camera pixels. The DMD pixels and those of the camera are found to be shifted. To correct the tilt, a pattern of intensity peaks is applied on the DMD. (a) The resulting beam is sent to a uniform fluorescence layer and imaged on the camera. (b-c) The tilt of the system is calibrated and the original pattern is thus corrected by a custom-made MATLAB code (d).*

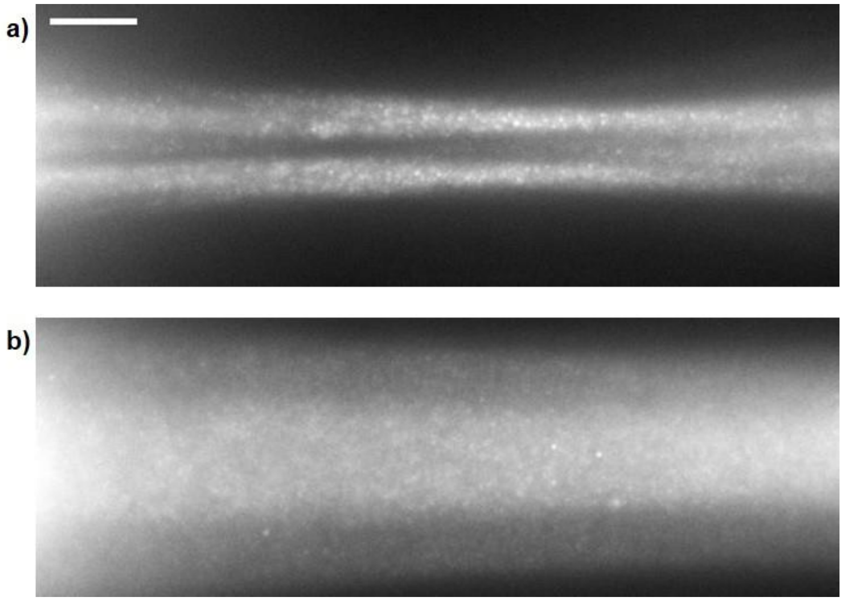


***Supplementary Figure 8 :*** *The double arc beam propagation. (a) The double arc beam as observed (x-y view) in the soSPIM chamber without scanning. The beam was sent through the chamber auto fluorescent polymer after reflection on the microfabricated mirror. (b) A x-y view of the double arc beam when scanned in the soSPIM sample holder.*

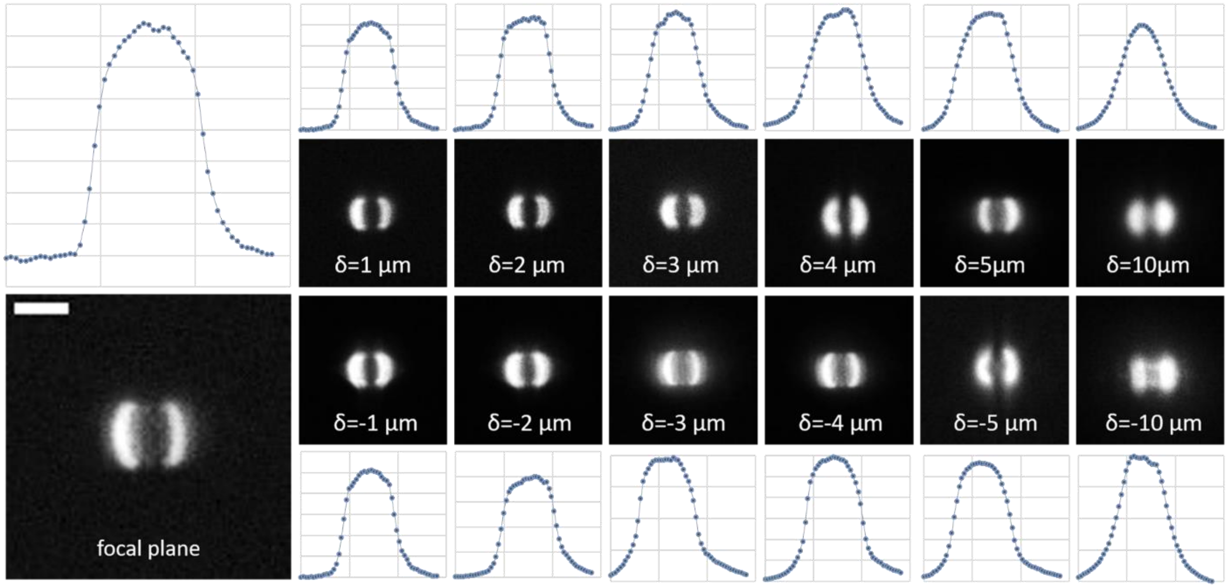


***Supplementary Figure 9 :*** *Lateral view of the double arc beam when propagating and the corresponding axial intensity profile: A uniform fluorescent layer was imaged when excited with the double arc beam without scanning. To simulate beam propagation, the microscope objective was fixed and the fluorescent layer was displaced axially by $\delta$. As such, the thin fluorescent layer intersects the beam at a distance $\delta$ from the focal of the objective. To observe the corresponding image at each displacement, the camera was defocused by a factor $\delta \times M^2$. The axial uniformity is preserved over $10\mu m$ propagation around the focal plane.*

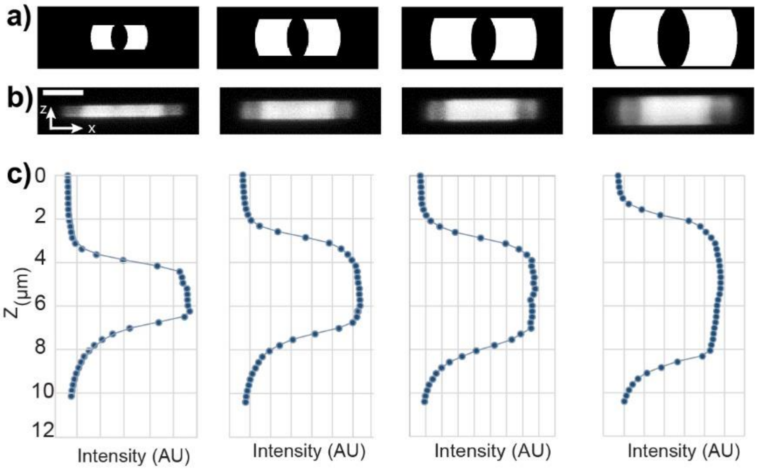


***Supplementary Figure 10 :*** *Uniformity of the axial intensity profile of the double arc beam for different light sheet thicknesses. (a) the applied double arc pattern on the DMD. (b) the shaped beam is scanned to produce a uniform axial beam along the z axis as observed by imaging a uniform fluorescence layer. (c) the resulting axial intensity profile. To note that the observed intensity does not reflect the exact intensity profile on the sample. In fact, the obtained images are the convolution of the intensity profiles at the sample and the detection PSF which tends to enlarge the intensity profiles.*

### Rayleigh length of the double-arc beam compared to a focused Gaussian beam

The width of a focused Gaussian beam of a wavelength $\lambda$, at a distance $z$ away from the position of waist is defined by

$$\omega(z) = \omega_0 \sqrt{1 + \left(\frac{z}{z_r}\right)^2}$$

$\omega_0$ is the waist size at the focus and $z_r = \pi \frac{\omega_0^2}{\lambda}$ is the Rayleigh length defined as the distance at which the width $\omega(z) = \omega_0\sqrt{2}$. The Rayleigh length corresponds to the distance over which the light sheet excitation remains confined and thus defines the range of the field of view that is useful for imaging.

For a light sheet obtained by 2D scanning of a focused Gaussian beam, the excitation beam is the superposition of multiple focused beams slightly displaced laterally. The waist along the optical axis can thus be defined as a $N$ times multiple of the waist $\omega_0$ of a single focused Gaussian beam ($N\omega_0$). As an approximation we consider the beam width as defined for a single beam $\omega(z) = \omega_0\sqrt{1 + \left(\frac{z}{z_r}\right)^2}$. This approximation stands as an underestimation of the width of the light sheet. Assuming an effective waist of $N\omega_0$, the effective Rayleigh length, at which the width is $\omega(z_{r0}) = \sqrt{2}N\omega_0$, is found for a distance $z_{r0} = z_r\sqrt{2N^2 - 1}$.

For a double arc shaped beam, the Rayleigh length is defined as for a Gaussian beam by $z_{r1} = \pi \frac{\omega\prime_0^2}{\lambda}$, where $\omega_0' = N\omega_0$. It can be rewritten $z_{r1} = z_r N^2$.

Comparing the Rayleigh length of both excitation modalities, the double arc shaped beam was found to extend over $N^2/\sqrt{2N^2 - 1}$ larger Rayleigh length compared to a light sheet of the same waist $N\omega_0$ obtained by 2D scanning of a focused Gaussian beam.

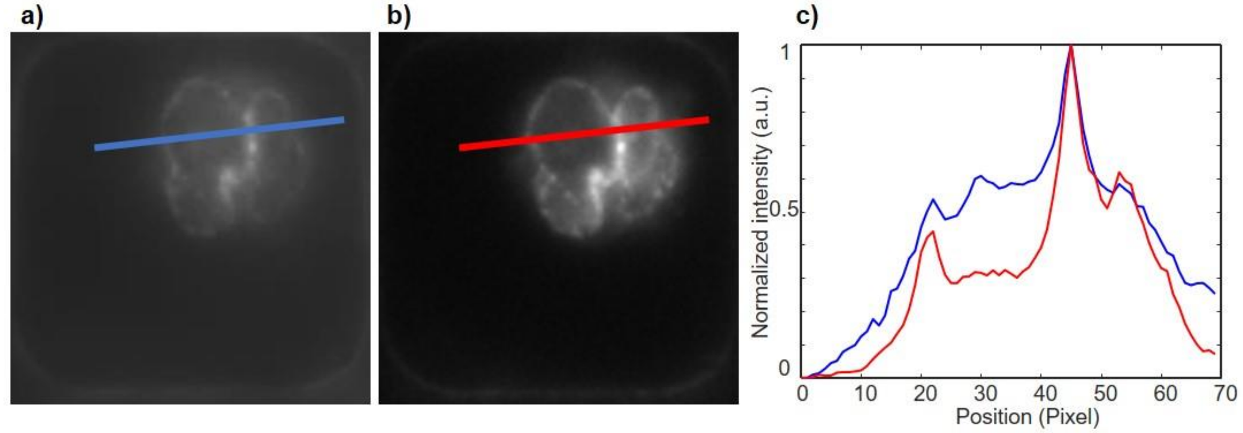


***Supplementary Figure 11 :*** *Signal to Noise Ratio of labeled nuclear proteins comparing widefield excitation (a) and V-LSFM (b). The two profiles across the nucleus are plotted on the same graph (c) following the normalization of the intensity between 0 and 1.*

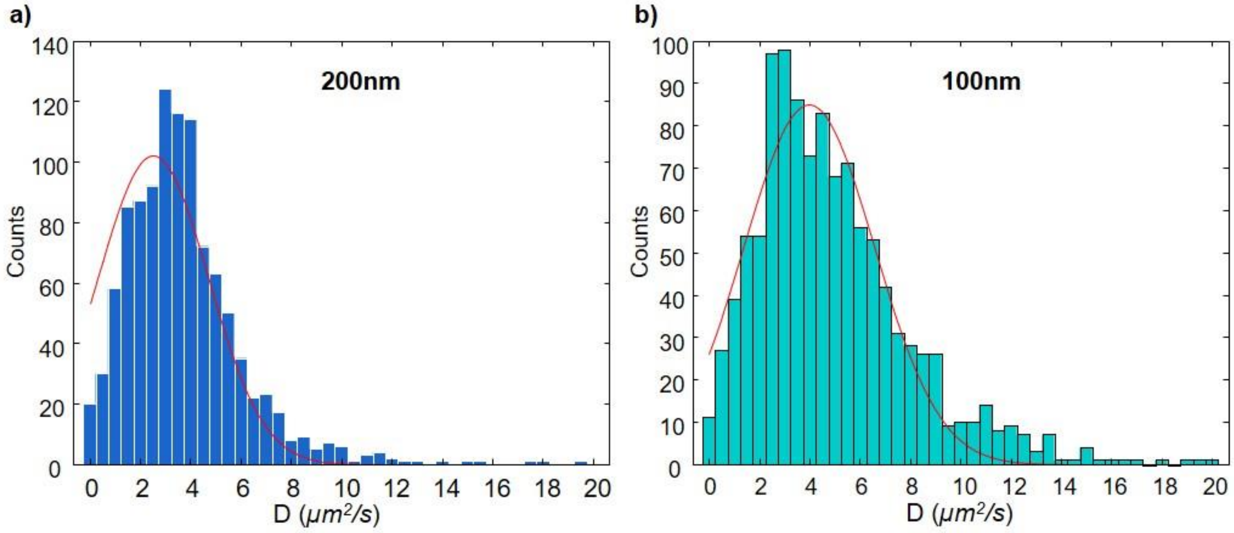


***Supplementary Figure 12 :*** *Histograms of 3D diffusion coefficients for (a) 200 and (b) 100nm particles diffusing in 3D and imaged with V-LSFM. The histograms are found to be distributed as a Gaussian centered around 2.5 and 3.9 $\mu m^2/s$ respectively. The measured values are in good agreement with the theoretical predictions of 2 and 4 $\mu m^2/s$. The difference and the width of the distribution can be explained by the inhomogeneous distribution of the size of the beads.*


### Supplementary videos:

**Supplementary video 1:** U2OS cells expressing NUP96 nuclear membrane protein fused ot GFP excited with a thin light sheet and imaged in MFM. The light sheet is scanned along the sample without displacing the focus or the sample.

**Supplementary video 2:** Raw data of beads diffusing in a solution and excited with a thick light sheet matching the observation volume of MFM. Fast diffusing beads tend to escape a focal plane after few frames.

**Supplementary video 3:** A reconstructed 3D trajectory of a bead illustrating the importance of fast imaging in recovering long trajectories

**Supplementary video 4:** Volumetric reconstruction of H2B proteins diffusing in the nucleus of 2 living U2OS cells. The volumetric data is overlaid with the reconstructed trajectories.

**Supplementary video 5:** Typical raw data acquired using V-LSFM showing single molecule Cohesin labeled with JF549 diffusing in live STEM cells. Images are acquired with 30ms exposure time a volume.
